# Mitotic activity shapes stage-specific histone modification profiles during Xenopus embryogenesis

**DOI:** 10.1101/2020.08.04.200550

**Authors:** Daniil Pokrovsky, Ignasi Forné, Tobias Straub, Axel Imhof, Ralph A.W. Rupp

## Abstract

Forming an embryo from a zygote poses an apparent conflict for epigenetic regulation. On one hand, the de novo induction of cell fate identities requires the establishment and subsequent maintenance of epigenetic information to harnish developmental gene expression. On the other hand, the embryo depends on cell proliferation, and every round of DNA replication dilutes preexisting histone modifications by incorporation of new unmodified histones into chromatin. Here we investigated the possible relationship between the propagation of epigenetic information and the developmental cell proliferation during Xenopus embryogenesis. We systemically inhibited cell proliferation during the G1/S-transition in gastrula embryos and followed their development until the tadpole stage. Comparing wild-type and cell cycle-arrested embryos, we show that the inhibition of cell proliferation is principally compatible with embryo survival and cellular differentiation. In parallel, we quantified by mass spectrometry the abundance of a large set of histone modification states, which reflects the developmental maturation of the embryonic epigenome. The arrested embryos developed abnormal stage-specific histone modification profiles, in which transcriptionally repressive histone marks were overrepresented. Embryos released from the cell cycle block during neurulation reverted back towards normality on morphological, molecular and epigenetic levels. These results indicate that replicational dilution of histone marks has a strong impact on developmental chromatin maturation. We propose that this influence is strong enough to control developmental decisions, specifically in cell populations that switch between resting and proliferating states such as stem cells.

## Introduction

Different cell types are distinguished by specific chromatin states, which help to target DNA-binding factors to specific genomic regions. Covalent post-translational modifications of histones (PTMs) contribute to these states in a pivotal manner (Lawrence, Daujat and Schneider, 2016). The most abundant and functionally studied histone modifications are acetylation, phosphorylation, and methylation, although many other modifications have been reported (Kebede, Schneider and Daujat, 2015). Transcriptionally active chromatin domains are characterized by a distinct array of histone marks. H3K27ac and H3K4me1 are associated with active enhancers (Creyghton et al., 2010), while high levels of H3K4me2/3 are found at the promoters of active genes (Barrera et al., 2008). Transcribed gene bodies are enriched in H3 and H4 acetylation (Myers et al., 2001), H3K36me3 (Pokholok et al., 2005), and H3K79me3 (Ng et al., 2003). In contrast, methylation of lysine residues 9 and 27 of H3 are hallmarks of repressive chromatin at silent gene loci (Alekseyenko et al., 2014; Ferrari et al., 2014). H3K27me3 is associated with the formation of facultative heterochromatin, whereas H3K9me2/3 has important roles in the formation of constitutive heterochromatin. Methyl marks on these two Lysines take part in regulating gene expression during development (Silva et al., 2003; Bracken et al., 2006). Altogether, histone PTMs ensure genomic integrity and control both adaptive and stable transcription modes to accommodate cell differentiation and physiological needs during development.

Histone modifications convey important information to early developmental programs in mammals (Jambhekar, Dhall and Shi, 2020). The regulation of epigenetic plasticity in mammals has been difficult to address in vivo so far, and our knowledge is largely derived from in vitro differentiation of pluripotent embryonic stem (ES) cells (Hajkova et al., 2008; Hemberger, Dean and Reik, 2009; Rugg-Gunn et al., 2010). The mechanisms, by which inductive signals shape the epigenome during ES cell differentiation, are still poorly understood. Xenopus laevis is a non-mammalian vertebrate model organism that allows to investigate in vivo the transition from pluripotency to committed cell states.

The epigenetic landscape of Xenopus embryos develops largely from an unprogrammed state, although maternally encoded proteins and epigenetic marks influence early transcriptional programs (Blythe et al., 2010; Hontelez et al., 2015). After maternal to zygotic transition (MBT) embryonic chromatin is considered permissive and receptive to inductive signals (Bogdanović, van Heeringen and Veenstra, 2012). Hierarchical clustering of histone PTMs has identified histone modification landscapes, which distinguish different developmental stages. We have termed these patterns as stage-specific histone modification profiles (HMP). They derive from quantitative fluctuations of histone PTMs, occurring during key developmental events such as germ layer formation, cell fate commitment and organogenesis. After the blastula stage HMPs gradually shift towards repressive histone PTMs (Schneider et al., 2011), in agreement with chromatin results from differentiating mammalian ES cells (Probst and Almouzni, 2008).

We have interpreted these stage-specific HMPs as part of a developmental program, which acts in parallel to the unfolding genetic pathways. Such a program might be controlled through mechanisms, which either modulate the expression or the kinetic activity of histone modifying enzymes. Cell proliferation could represent an additional mechanism, since histone incorporation during S-phase dilutes preexisting histone modifications and both cell cycle length and proliferative status of embryonic cells are subject to dramatic changes. At early stages of Xenopus development (similar to zebrafish and Drosophila), the cell cycle is driven by an autonomous biochemical oscillator, which is unaffected by developmental signals or checkpoints (Newport and Kirschner, 1982; Kimelman, Kirschner and Scherson, 1987; Anderson et al., 2017). Cleavage divisions are extremely rapid (from 10 to 30 min) and lack gap phases. After MBT, gap phases appear, the cell cycle lengthens, and becomes asynchronous (Desnitskiy, 2018). Subsequently, an increasing proportion of cells exits the cell cycle during differentiation. Indeed, the first postmitotic cells appear already at the early neurula stage (Hartenstein, 1989). They are thought to be in G0 phase; however, it was recently shown that quiescent cells can also be found in G2 (Sabherwal et al., 2014). This developmental regulation of cell division outside of the G1 phase allows to quickly reinitiate cell divisions later in development. The finding, that in both Xenopus and Drosophila neurogenesis periods of intense proliferation are interrupted by phases of quiescence, supports this notion of the cell cycle being used in a regulatory manner (Thuret, Auger and Papalopulu, 2015; Ramon-Cañellas, Peterson and Morante, 2019). These changes in the cell cycle are accompanied by changes in local chromatin structure. Important histone modifications such as trimethylated H3K27 or H4K20 are not present at significant levels prior to zygotic genome activation, but greatly increase after the MBT, when large-scale changes in chromatin modifications happen (Vastenhouw et al., 2010; Lindeman et al., 2011). Together, this data indicates that major changes in cell cycle regulation and the appearance of stage-specific HMPs occur in a developmentally coordinated manner.

Cell division can be blocked in Xenopus from gastrulation onwards by incubating embryos with the DNA synthesis inhibitors Hydroxyurea and Aphidicholin (“HUA”) (Harris and Hartenstein, 1991). Here we demonstrate that Xenopus embryos, arrested systemically during the G1/S transition during gastrulation (NF10.5), develop largely normal and in synchrony with control siblings until the tadpole stage (NF32). However, cell cycle-arrested embryos display an altered histone modification landscape, in which specific modifications accumulate both precociously and in excess. If arrested embryos are released back into proliferation, the chromatin landscape reverts towards normal. Thus, our data establish that stage-specific HMPs are sensitive to changes in mitotic activity.

## Results

### HUA treatment: experimental design

Early Xenopus embryos rely on the consumption of maternally supplied molecules, which restricts the available repertoire of cell cycle manipulations. We have chosen a systemic cell cycle arrest, achieved by a combination of two small-molecule inhibitors: Hydroxyurea and Aphidicolin (from here on called HUA condition). When applied to cells in culture, both drugs effectively block the initiation of DNA replication and consequently cell division, without obvious side effects on cell viability or differentiation capacity (Jensen, 1987; Maurer-Schultze, Siebert and Bassukas, 1988). This effectively leads to a cell cycle arrest at the G1/S transition. Harris and Hartenstein have used these inhibitors to demonstrate that cell division is dispensable for neural induction, neuronal differentiation, and for neural tube formation (Harris and Hartenstein, 1991). We have confirmed their finding that whereas HUA treatment before gastrulation (i.e. at NF9) leads to aborted development, it is principally compatible with embryonic development until tadpole stages when started shortly after gastrulation onset (NF10.5) (for more information see Method section and Fig S1A).

In this work we have assessed the effects of the systemic cell cycle arrest by two types of experiments (Fig 1). Experiment type A (“permanent arrest”) implies a continuous HUA or Mock treatment on sibling embryos from NF10.5 until NF32. Mock treatment represents incubation in 2% DMSO, which is the solvent for Aphidicolin. Experiment type B (“transient arrest”) consists of continuous HUA and Mock treatments plus an additional HUA washout (HUAwo) condition. In the latter case, embryos were temporarily incubated with HUA from early (NF10.5) to late (NF13) gastrula stages, and then returned to Mock solution. Experiment type B tests the reversibility of the cell cycle arrest and its developmental consequences.

**Figure 1.**
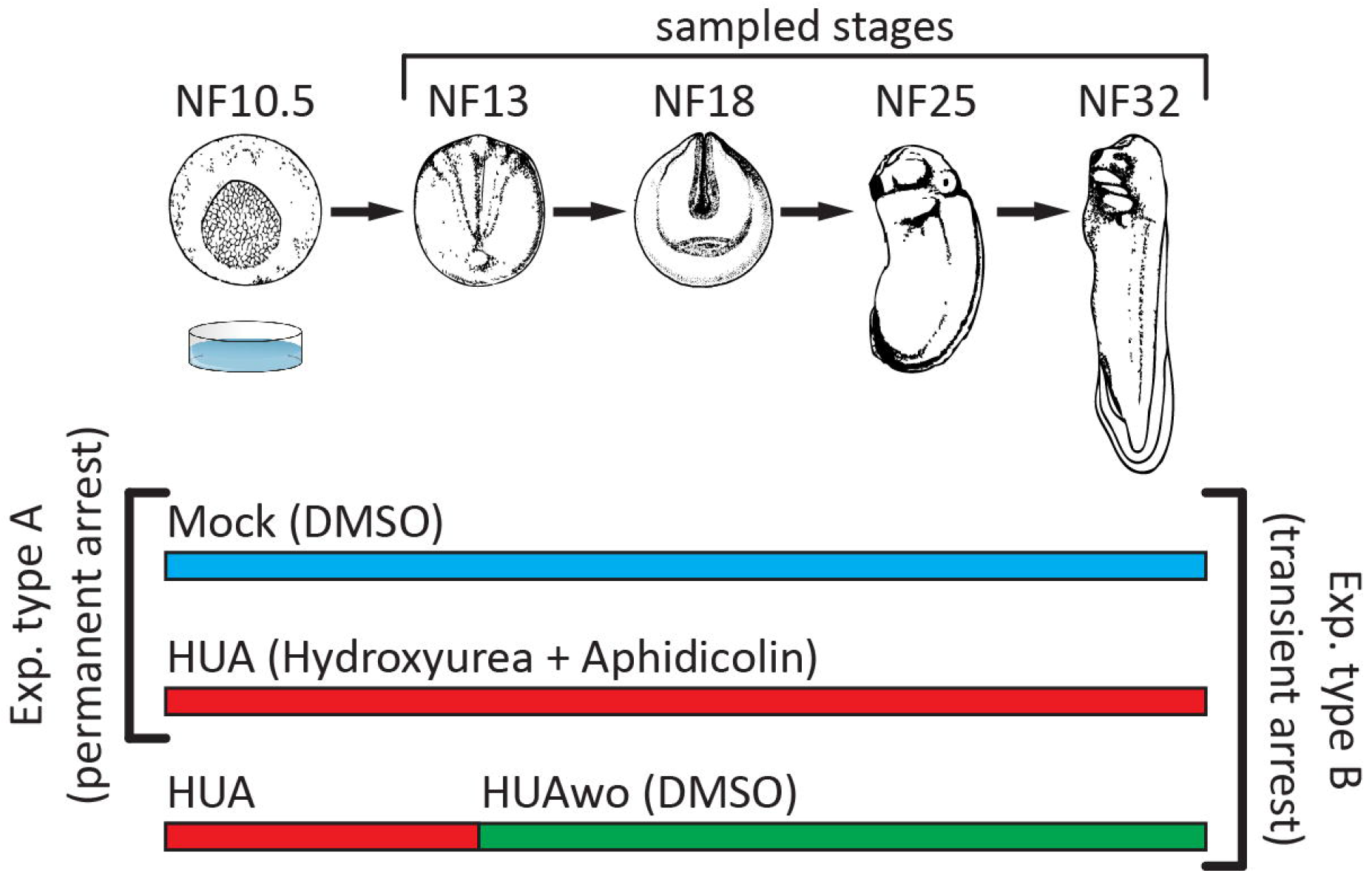
Experimental setup. Top – A schematic representation of Xenopus laevis embryonic development. Developmental stages (NF) according to Nieuwkoop and Faber, 1994. Stages used for mass spectrometry of histone modifications and embryonic analyses are characterized by the following features: NF13 – late Gastrula, germ layer specified; NF18 – Neurula, germ layer patterning and differentiation; NF25 – Tail bud stage, organogenesis; NF32 – early Tadpole, body plan established. Bottom part – we perform two types of experiments: Experiment type A (G1/S block) – embryos are split in two groups, which from NF10.5 on are continuously incubated in HUA solution (Hydroxyurea and Aphidicolin) or control solution (DMSO as carrier). Experiment type B (transient arrest) – embryos are split in three groups – continuous Mock, continuous HUA and transient HUA. In the last group, HUA solution is replaced at NF13 with DMSO solution.

For all experiments embryo siblings were collected at four developmental stages: NF13 (early neurula), NF18 (late neurula), NF25 (tail bud) and NF32 (tadpole) (Fig 1). By the early neurula stage, germ layers and body axes have been determined. The embryonic patterning increases the cellular diversity of the embryos during neurulation. At the tailbud stage, the first differentiated tissues such as skeletal muscle and the mucociliary epithelium of the larval skin have been formed. The end point of the analysis (NF32) lies behind the phylotypic stage of Xenopus (NF28-31), when evolutionary conserved gene expression programs have established the vertebrate body plan (Irie, Satoh and Kuratani, 2018).

### HUA effects on embryo viability and cell proliferation

At the first sampling stage (NF13), there was no difference in viability between proliferating controls and HUA arrested embryos (Fig S1A). As reported before (Harris and Hartenstein, 1991), HUA treatment reduced embryonic viability after NF18, such that the survival rate was about half of that of Mock embryos at NF25 and NF32. Staining for activated Caspase 3 indicated that both Mock and HUA treated embryos contained apoptotic cells in a variable, but comparable extent (Fig S2). Depending on individual egg batches, arrested embryos survived up to stage NF41.

To test the efficacy of the HUA incubation we stained embryos for the histone H3 Serine 10 phospho mark (Fig 2), which accumulates in M-phase cells (Wang and Higgins, 2013). We determined the number of H3S10Ph-positive cells from the entire surface of the embryo with Image J (Fiji). Already at NF13, HUA embryos contained more than 8-fold less mitotic cells than Mock embryos, a difference that further increased to 17-fold by NF32. We conclude that S-phase dilution of chromatin marks is efficiently prevented by HUA treatment.

**Figure 2.**
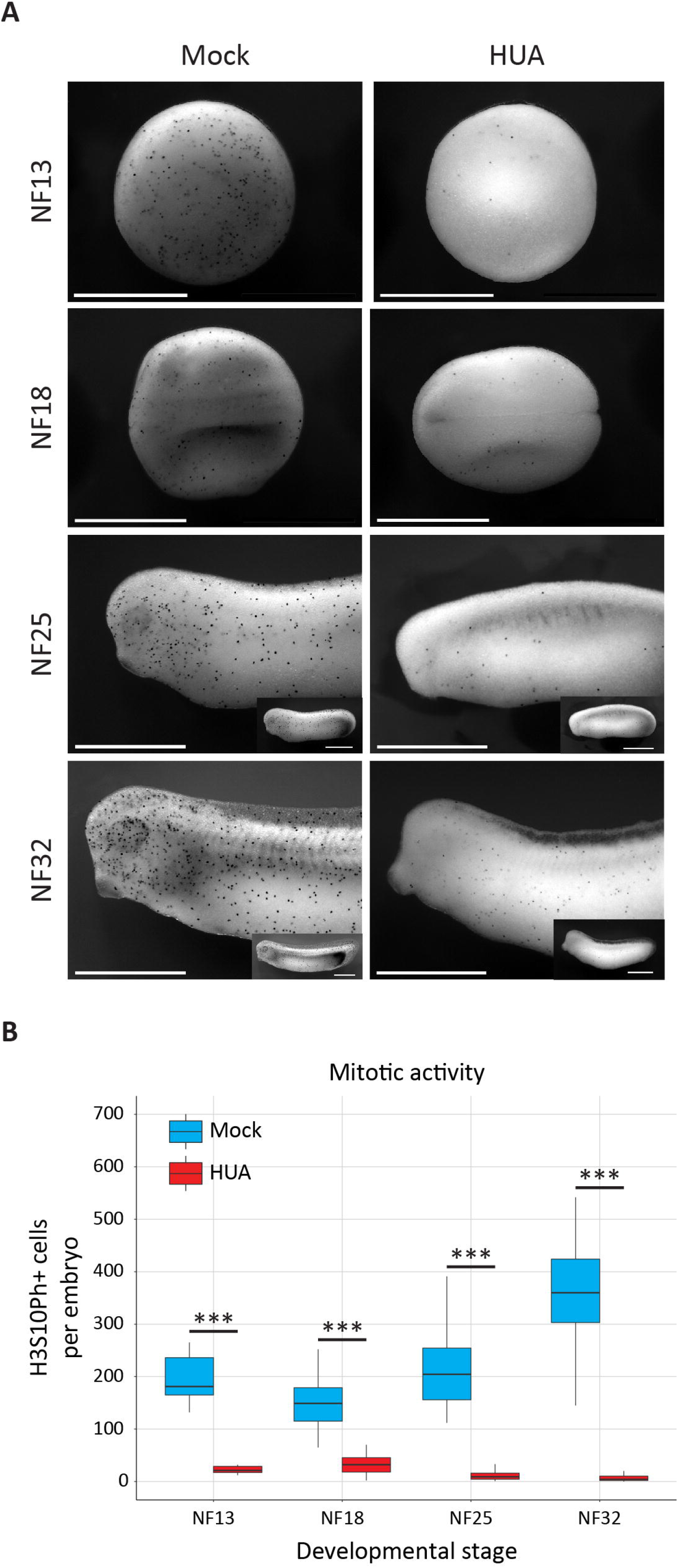
Continuous HUA treatment inhibits mitotic activity from gastrula to tadpole stages. A) Immunocytochemical staining (ICC) for the mitotic histone mark H3S10Ph. Black dots represent mitotic cells. Elongated, older embryos are recorded as anterior halves, i.e. at same magnification as younger stages, and in whole mount views as inserts. Scale bars: 1mm. B) Abundance of mitotic cells in Mock and HUA treated embryos. Box plots based on H3S10Ph-positive cells present on the recorded surface of embryos (n=3 biological replicates/condition; Student’s t-test [unpaired, two-tailed]; *** p<0.001).

### HUA effects on morphogenesis

The cell cycle arrested embryos completed gastrulation in synchrony with Mock controls (Fig 3A). From mid-neurula onwards (stage NF18/19) the closure of the neural tube was delayed, consistent with previous findings (Harris and Hartenstein, 1991) (Fig S1B). At later stages, HUA embryos lacked a postanal tail, melanocytes, and the eye Anlage remained rudimentary. Note that the size of arrested and Mock treated embryos was almost the same until stage NF25 (Fig 3A). Since holoblastic cleaving Xenopus embryos rely completely on maternal storage, cell proliferation subdivides the zygote into smaller and smaller cell progeny, while maintaining the maternally derived mass. Consistent with the observed mitotic arrest we confirmed that HUA embryos consisted of bigger cells at the neurula stage (Fig S1B). Confocal imaging indicates that epidermal cells of HUA embryos were clearly spread out over a larger area than proliferating cells of control embryos, and their nuclei appear enlarged (Fig 3B). H3S10Ph-positive cells were detected in Mock controls, but not in HUA embryos (Video S1 and Video S2). Overall, these observations document a remarkable compensation on the cellular level, which enables HUA embryos to undergo morphogenesis in apparent synchrony with control siblings.

**Figure 3.**
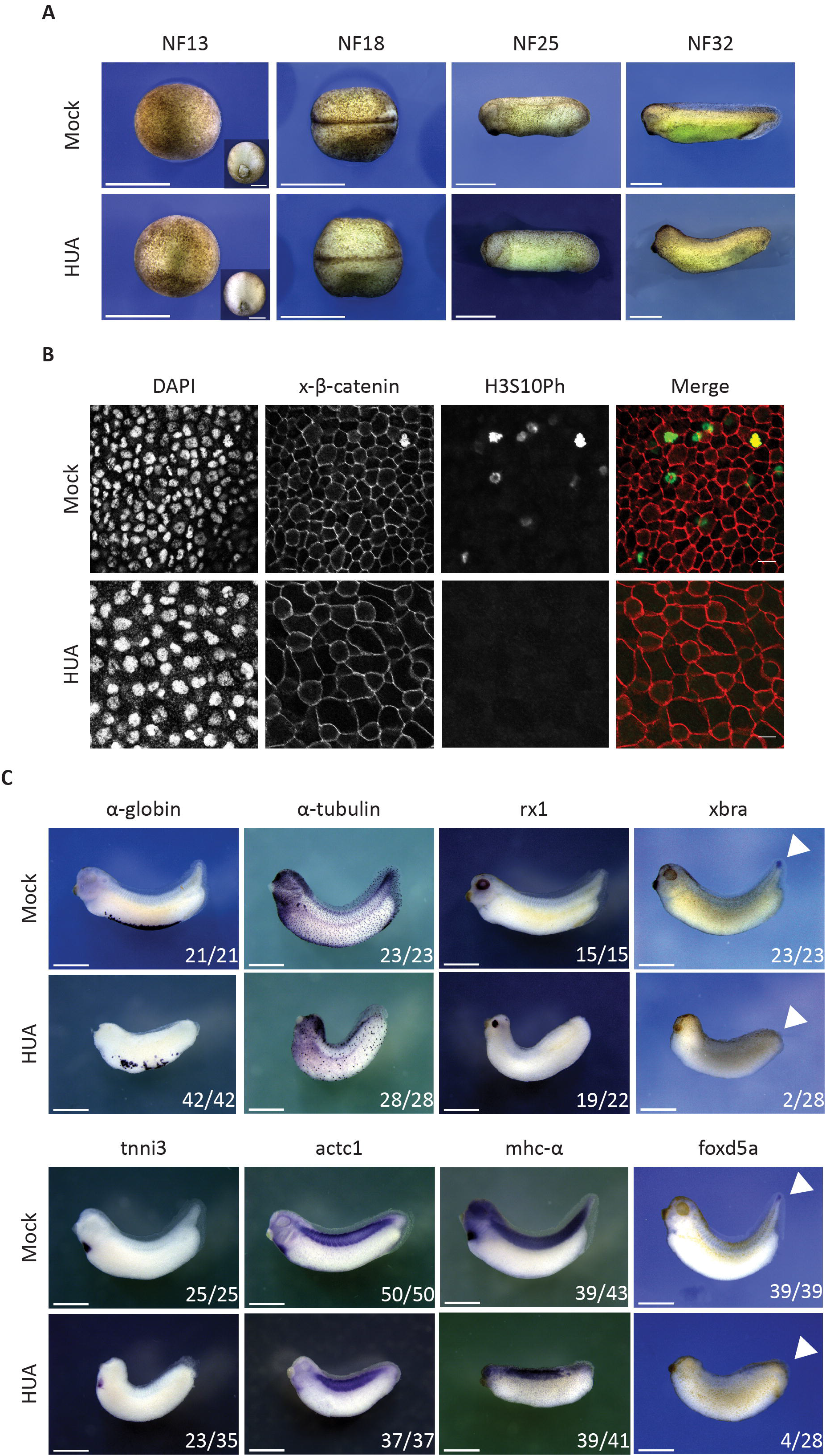
Morphological development of HUA-arrested embryos. A) During gastrulation, Mock and HUA embryos are indistinguishable. At NF18, the latter show a delay in neural tube closure. More severe malformations are detectable at stages NF25 and NF32, most notably reduced eye formation and absence of tail bud. Scale bars: 1mm. B) Cell size at Tail bud stage. Flattened Z-stack images show fields from embryonic skin at constant magnification (scale bars: 20μm). Immunofluorescence detects cell borders (beta-catenin), nuclei (DAPI), mitotic cells (H3S10Ph). GIFs from Z-stacks are presented in the supplementary material (Video S1 and S2). C) Whole mount RNA in situ hybridization for indicated marker genes. Images are representative for the majority of Mock or HUA treated embryos from 3 biological experiments. While mRNAs of α-tubulin and rx1 (skin, neuronal), α-globin, tnni3, actc1 and mhc-α (mesodermal) are detected in both conditions, xbra and foxd5a (tail blastemal) are absent in HUA embryos. Numbers indicate embryos positive for the marker over the total number of analyzed embryos. Scale bars: 1mm.

### Differentiation of cell cycle-arrested embryos

Cell proliferation and differentiation are in most cases mutually exclusive processes (Weintraub et al., 1991). To investigate the developmental relationship of Mock and HUA-treated embryos, we investigated the tissue-specific expression of a panel of marker genes (Fig 3C, Table S1). Whole mount RNA in situ hybridisation revealed that most of these markers (17 out of 19) were expressed at the correct site, although in smaller domains. Only two genes were not transcribed in the majority of HUA-treated embryos. Xbra and foxd5 are normally expressed in the tail, which HUA-embryos fail to form. To rule out that the absence of these mRNAs in cell cycle arrested embryos was due to a developmental delay, we compared by qRT/PCR the timing and relative levels of gene transcription between Mock and HUA treated embryos (Fig S1C). We examined 6 genes, which become induced at different time points – pax6 and actc1 at gastrulation, twist and myt1 during neurulation, and tnni3 and fabp2 in tadpoles – and normalized the relative mRNA levels to the housekeeping gene odc. A significant difference in gene expression between Mock and HUA embryos was found only for pax6, which had disappeared by stage NF32. The mRNA levels of the other 5 genes, including the late-induced fabp2 gene, were proportionate between the two conditions. This indicates that genes were activated around the proper time, arguing against a general delay in the development of HUA embryos.

Although HUA-treated tadpoles were morphologically impaired, they responded to touch stimulation by muscle twitching, and at least some of them were capable of mounting a flight response (Video S3 and Video S4). This burst of swimming activity involves a physiological connection between sensory neurons, the CNS, and the body wall musculature (Roberts, Li and Soffe, 2010).

The morphological and molecular analyses demonstrated that embryonic development proceeds in the absence of cell proliferation, although the Anlagen of some organs like tail, fin and eyes were clearly compromised. These findings are in agreement with and extend previous observations on HUA-treated embryos (Harris and Hartenstein, 1991). We conclude that blocking the cell cycle from gastrula stages onwards is compatible with largely normal and apparently synchronous differentiation of the embryo.

### Stage-specific histone modification profiles in HUA embryos

To investigate the impact of the cell cycle arrest on embryonic chromatin, we compared the posttranslational histone modification profiles in proliferating and HUA arrested embryos by quantitative mass spectrometry (for more information see Method section). In total, this study quantified 65 histone modification states across 4 developmental stages in 2 experimental conditions, each in 3 biological replicates. The combination of all modification states at a certain developmental timepoint constitutes the so-called stage-specific histone modification profile (HMP) (Schneider et al., 2011). To visualize differences in the HMPs between cell cycle-arrested and control embryonic chromatin, we performed hierarchical clustering analysis with all 24 samples to build a single, stagespecific heatmap (Fig 4A). This clustering was based on the absolute intensity values of endogenous histone peptides.

The first three levels of the dendrogram on the left axis of the heatmap revealed five clusters with the following features. Cluster 1 grouped histone PTMs, which undulate in control embryos, i.e. they were more abundant at stages NF13 and NF25 than at stages NF18 and 32. In cell cycle-arrested embryos, this pattern collapsed into a single high point at NF25. Cluster 2 included modification states, which gradually increased their abundance under both experimental conditions. Cluster 3 represented modifications, which were more abundant in HUA-arrested chromatin compared to control chromatin, whereas in clusters 4 and 5 highly abundant modification states in Mock embryos were downregulated in HUA samples.

Despite the fact that the cell cycle arrest alters the abundance of nearly every analysed modification to varying extent, HMPs are clearly discernible for both conditions (Fig 4A). To obtain further insight into their difference, we performed PCA analysis for the whole data set (Fig S3). The two conditions are partly separating, particularly clear for late gastrula control and tadpole HUA samples. In general, chromatin features of younger HUA samples seem to co-segregate with those of older controls.

### HUA effects on individual histone marks

The global, absolute heatmap indicated that the HUA block resulted in a different developmental HPM profile. To obtain a better understanding of how the histone modification states differed between the two conditions, we converted the absolute abundance of individual histone modifications into relative proportions for those modification states, which are located on a single tryptic histone peptide (Fig S4, and Table S5). We then generated a second heatmap, to display the fold-change between Mock and HUA conditions at a given stage (Fig 4B).

The relative ratio heatmap supports the permanent nature of the G1/S cell cycle arrest by revealing a more than 10-fold reduction of the mitotic chromatin mark H3S10Ph. This data confirmed that HUA treatment affected not only superficial cell layers, but all embryonic tissues. Furthermore, the overall levels of H3K4me2/me3 were maintained in HUA embryos at all four stages, suggesting that the number of active and poised promoters can be maintained in non proliferating embryos, consistent with the largely correct expression of cell-type specific markers (Fig 4B and Fig 3C). Other modifications deviated significantly between the two conditions. The embryonic epigenome of HUA-treated embryos contained strongly reduced levels for some modifications, notably the acetylated states of K9 and K27 on histone H3. These modifications are thought to prevent the writing of repressive methylation marks on chromatin harbouring cis-regulatory DNA elements (Sabbattini et al., 2014; Zhang, Cooper and Brockdorff, 2015), a prerequisite for de novo enhancer activation (Smith and Shilatifard, 2014). Beyond the H3S10P mark, the next largest differences in HUA-arrested chromatin were found for the trimethylated states of Lysines 9 and 27 on H3, and Lysine 20 on H4. These sites became globally elevated by more than 2,5-fold compared to the chromatin of control embryos. While higher levels of H3K27me3 and H4K20me2/3 were anticipated, since their levels are linked to cell cycle progression (Alabert et al., 2015), the increase in H3K9me3 suggests that the HUA cell-cycle arrest occurs partly in S-phase (Di Micco et al., 2011).

Altogether, we demonstrated that incubating Xenopus embryos in HUA solution was an efficient way to systemically block cell proliferation. The permanent block had consequences on defined morphological aspects of development, but altered also stage-specific histone modification profiles. Most notably, this had an impact on specific histone modifications known to be involved in epigenetic memory and enhancer activity (Lee and Mahadevan, 2009; Smith and Shilatifard, 2014). The chromatin differences between HUA and control embryos indicate that the cell cycle state — possibly cell proliferation as such — coordinates the developmental decoration of chromatin with covalent histone modifications.

### HUA effects on embryogenesis are reversible

The observed chromatin changes in response to the HUA treatment could represent a non-physiological derailment, rather than a kinetic consequence of the G1/S arrest. To distinguish between these possibilities, we performed a transient arrest (HUAwo) (Fig 1).

Only four hours after returning to Mock condition (i.e. at midneurula stage – NF18), H3S10Ph-positive cell numbers were significantly increased compared to siblings maintained in HUA (Fig 5A, and Fig S5). At the endpoint of the experiment, mitotic cells were equally abundant in HUAwo and Mock conditions, indicating full recovery of the mitotic activity within 13 hours after inhibitor removal. In addition, the survival rate of HUAwo embryos increased, compared to continuously arrested embryos, and was similar to that of Mock embryos (Fig S1A).

**Figure 4.**
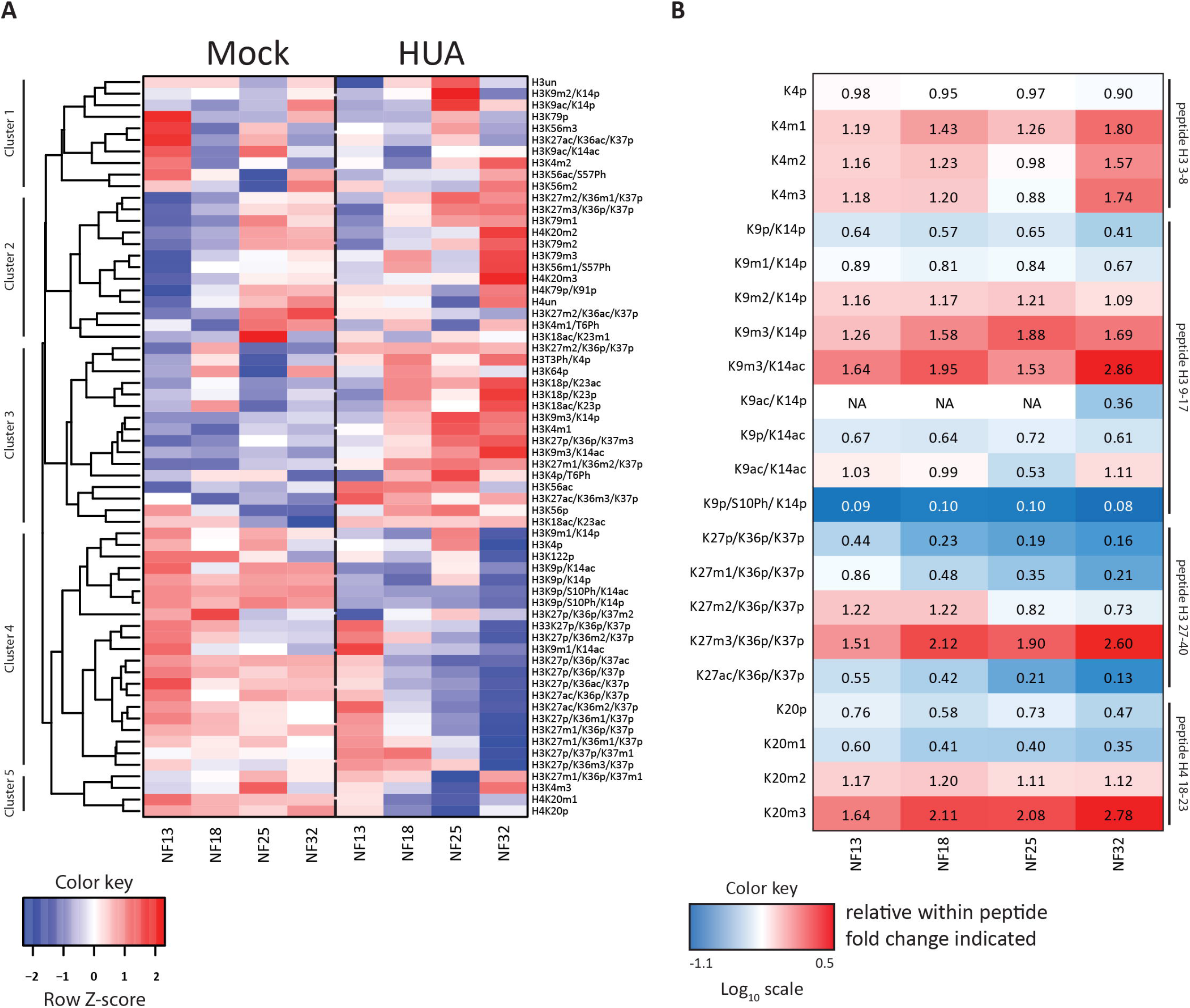
Mitotic activity shapes stage-specific histone modification profiles. A) Heatmap of the absolute abundances of individual histone modification states, normalized to the corresponding R10 spiketides. Data in columns represent an average from 3 biological replicates/condition. Color key: row Z-score. B) Ratio heatmap of relative histone modification abundances for selected histone marks. In the first step, relative distributions of the indicated modification states were calculated in percentages within each tryptic peptide (see Fig S4). Then, HUA values were divided by Mock values to produce the relative change for each histone modification state between the two conditions. Color key is based on the Log10 scale. Numerical values in the cells indicate the fold-change. Values greater 1.0 indicate an increase in HUA condition, values smaller 1.0 indicate a higher abundance in Mock condition. NA – not applicable, when values in Mock were 0.

Morphological hallmarks also recovered towards normality in HUAwo embryos (Fig 5B). Unlike continuously arrested embryos, the HUAwo group frequently developed postanal tails of nearly normal length, a clearly distinguishable fin surrounding trunk and tail. The recovery of the tail structure was accompanied by the recurrence of xbra and foxd5a mRNAs at the tail tip of all HUAwo embryos (Fig 5B).These findings indicate that the major morphological defects of the HUA arrest are reversible, at least when the inhibitors are removed during neurulation.

**Figure 5.**
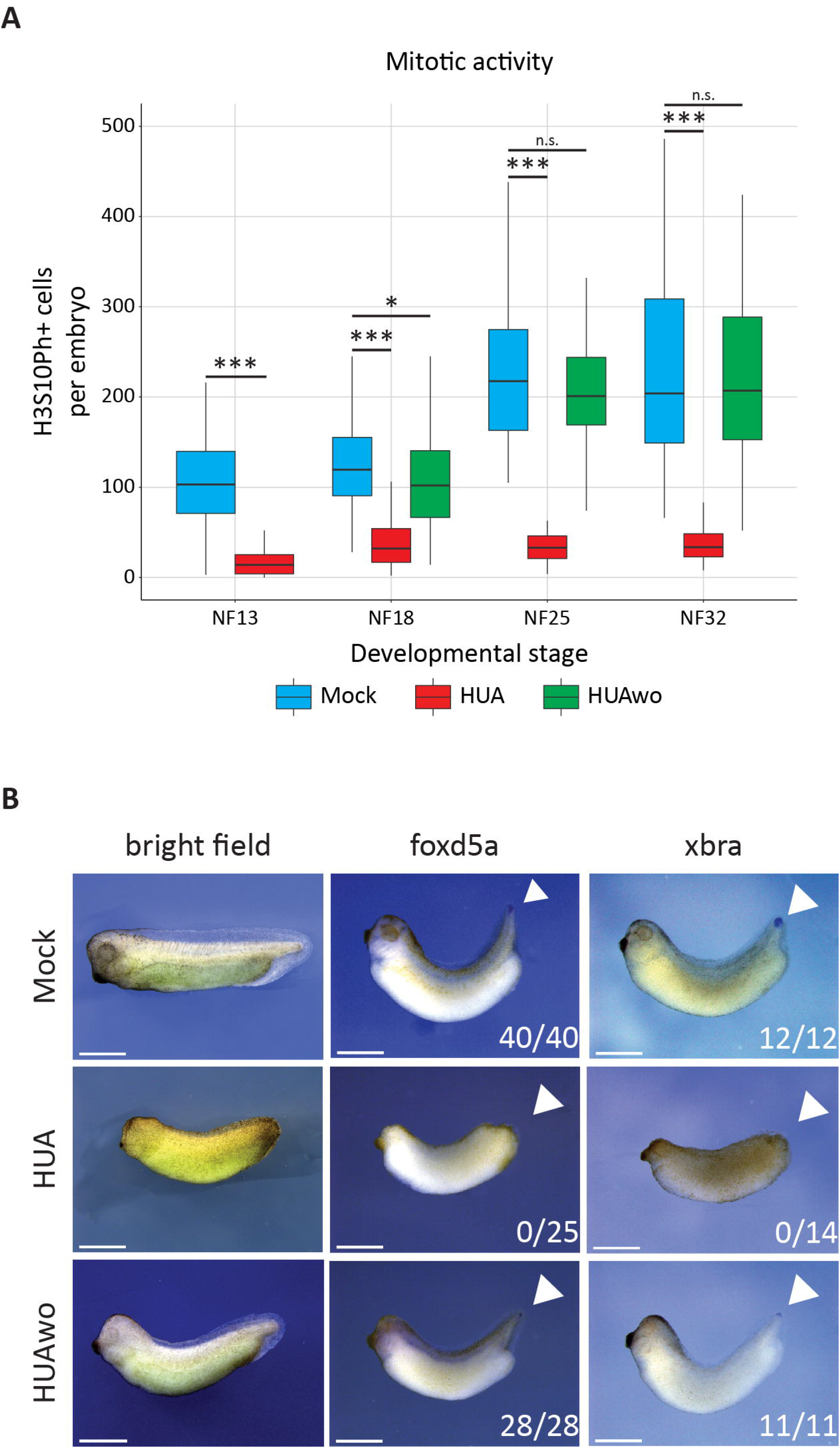
The HUA effects on embryogenesis are reversible. Embryos, which were transiently incubated in HUA solution and thus mitotically arrested, were returned to Mock solution at NF13. A) Abundance of mitotic cells in Mock, HUA treated, and HUA wash-out (HUAwo) embryos. Box plots based on H3S10Ph-positive cells present on the recorded surface of embryos, displayed in Fig S5 (n=3 biological replicates/condition; Student’s t-test [unpaired, two-tailed]; *** p<0.001; * p<0.05; n.s. – not significant). B) Morphological and molecular features of embryos at early tadpole stage. In contrast to continuous HUA treated embryos, HUAwo embryos regain eye cups, fin and tailbud structures. In addition, they express foxd5a and xbra mRNAs in the growth zone of the tailbud, comparable to Mock embryos. Numbers indicate embryos positive for the marker over the total number of analyzed embryos (n=3 biological replicates/condition). Scale bars: 1mm.

Encouraged by the results, we next performed mass spectrometry analysis on sibling cohorts of Mock, permanent HUA, and HUAwo embryos at tadpole stage (NF32). The results were visualized in a heatmap, based on relative values (Fig 6A, and Table S6). As for the experiment type A, the fold-change of functionally well annotated histone PTMs was calculated from their relative proportions (Fig 6B, and Fig S6). In total, almost three quarters of the modifications were closer to the levels found in control embryos than to HUA chromatin, although in most cases they do not reach wildtype abundance.

**Figure 6.**
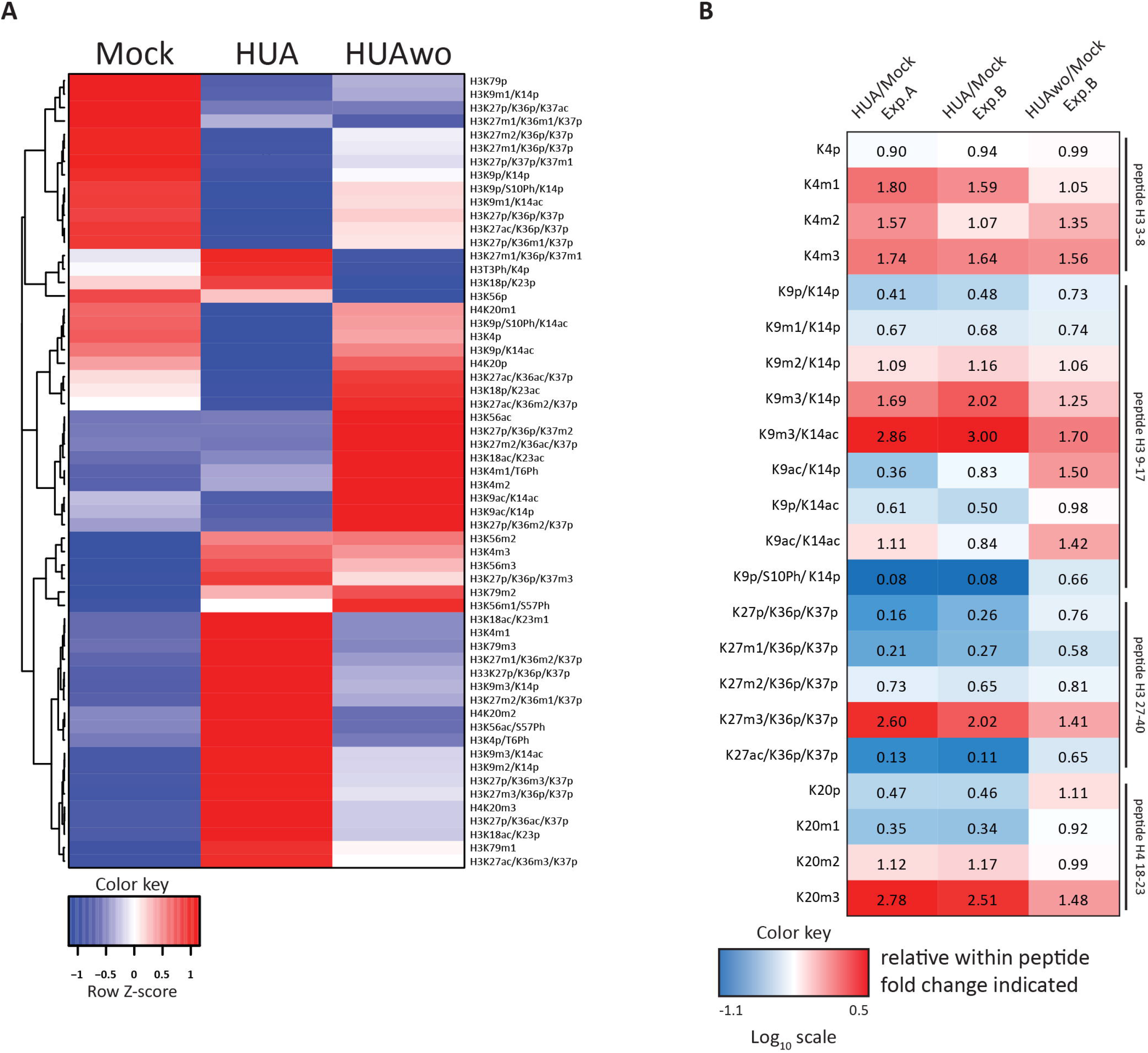
HUA effects on stage-specific histone modification profiles are reversible. A) Heatmap of the relative abundances of individual histone modification states under the indicated conditions. As before, the relative distributions of the indicated modification states are calculated in percentages within each tryptic peptide. Data in columns are collected from early tadpole stage embryos (n=1), independently from the data of Experiment type A. Note: relative differences between Mock and HUA samples of Experiment type B are highly similar to the results of Experiment type A (Fig S6). Color key, row Z-score. B) Ratio heatmap of relative histone modification abundancies for selected histone marks. First, the relative distributions of the indicated modification states are calculated as percentages within each tryptic peptide (Fig S6). Then, the relative values are presented as ratios between sample pairs indicated on top, showing the relative change for each histone modification state between the two conditions. Color key is based on the Log10 scale. Numerical values in the cells indicate the fold-change. Values greater 1.0 indicate an increase in HUA condition, values smaller 1.0 indicate a higher abundance in Mock condition. NA – not applicable, when values in Mock were 0. The first two columns illustrate the similarity between Experiment type A and B, the information in the third column shows the similarity between HUAwo and Mock samples.

In detail, we found that H3S10Ph levels were 8-fold higher in transiently arrested (66% of Mock) versus constantly arrested (8% of Mock) embryos. The return to proliferation is also reflected in increased H4K20me1 levels. In proliferating cells this modification reaches the maximum in G2- and M-phases, when PR-Set7 monomethylates new histones incorporated during the previous S-phase (Wu et al., 2010; Beck et al., 2012). The observed doubling in H4K20me1 levels (Fig 6B, and Fig S6) indicates that upon inhibitor washout, cells can go thrcough S-phase and mitosis. Also the H3K27me1 mark, which promotes the transcription of H3K36me3 decorated genes in ES cells (Ferrari et al., 2014), is recovering as well as the abundance of acetyl marks on H3K9 and H3K27. The latter marks protect histone H3 from the repressive influence of trimethylation at these Lysines, consistent with a transcriptionally more permissive state of chromatin. Indeed, levels of the repressive chromatin modifications H3K9me3, H3K27me2/3, and H4K20me3 were lower in the HUAwo condition, compared to permanently arrested embryos. Whether these changes are causally connected, i.e. acetylation marks replace methyl marks, or occur at independent chromatin regions, is not known. Nevertheless, the readjustments in the chromatin landscapes of embryos released from the G1/S block is in agreement with the observed morphological improvement.

Overall, these results document that HUA-arrested embryos can re-initiate cell proliferation, when the inhibitors are washed out at late gastrula. In contrast to embryos, treated continuously with the cell cycle inhibitors, the HUAwo embryos show a higher vitality with improved tissue formation and gene expression. These changes are accompanied by a partial restoration of their normal histone modification landscape. Taken together, the results of the inhibitor washout experiment testified to the non-toxic nature of the HUA-dependent cell cycle arrest in Xenopus embryos and substantiated the impact of cell proliferation on stage-specific histone modification profiles.

## Discussion

Establishing and maintaining the epigenetic information of covalent histone modifications faces a fundamental problem in proliferating cells, i.e. a two-fold replicational dilution of preexisting histone marks during S-phase needs to be matched by the biological kinetic rates of the enzymes decorating the chromatin landscape. While one might assume that evolution has ensured a robust balance between dilution and writing of histone PTMs, which adequately meets the physiological requirements of cells to acquire and maintain gene expression profiles and genome stability, recent work has indicated that this balance is delicate and could provide a resource for epigenetic regulation (Méchali, 2010; Tsubouchi and Fisher, 2013; Steffen and Ringrose, 2014; Jadhav et al., 2020).

In this study, we have addressed the impact of replicational dilution on the histone modification landscape during X. laevis embryogenesis, in which the cell cycle naturally undergoes massive changes. In this system, modulations in cell cycle phases, cell cycle length and differentiation-coupled withdrawal from cell cycle are tightly coordinated with developmental programs. We had previously noticed that Xenopus development is characterized by stage-specific fluctuations in steady state abundance of histone modifications, which we suspected to encode epigenetic information for the regulation of developmental processes (Schneider et al., 2011). These assumptions have been tested here by employing a small molecule inhibitor strategy (HUA), which had been pioneered by Harris and Hartenstein (1991). We have achieved a nearly complete arrest of cell proliferation from late gastrula stage on, which we have used to monitor the consequences for both the histone modification landscape and for embryonic differentiation by RNA in situ hybridization for marker genes, while embryos develop into tadpoles.

Importantly, HUA treatment is not tolerated by younger (blastula) embryos, which die within a few hours after inhibitor application. Therefore, we cannot investigate with this method the immediate seeding of histone marks in the wake of zygotic genome activation at MBT. This temporal restriction provides a plausible explanation for how HUA arrested embryos can finish the formation of germ layers and embryonic axes with the same efficiency as controls (Fig 3 and S1) – i.e. the cell cycle arrest occurs mainly after these events. With notable exceptions discussed below, this is reflected in the maintenance of many gene expression domains in HUA embryos (Fig 3).

While our analysis of the cellular composition and differentiated status of cells and organs is naturally limited, we note that HUA embryos at the tadpole stage are capable of muscle contraction, both spontaneously and in response to touch stimulation (Video S3 and Video S4). Furthermore, their epidermis contains multiciliated cells (see Fig 3C, alpha tubulin staining), which are fully differentiated at the tailbud stage. Finally, the HUAwo experiment demonstrated that morphological phenotypes, aberrant gene expression patterns, and the histone modification landscape are all largely recovering towards normality. We therefore conclude that changes in the histone modification landscape arise as a consequence of the G1/S-phase cell cycle arrest.

By far the largest differences in abundance were measured for the mitotic H3S10Ph mark, catalyzed by aurora kinases (Willems et al., 2018). Throughout the entire time course, the abundance of this G2/M mark was twelve-fold reduced in arrested embryos. Therefore, proliferating cells represent only a minor contamination in HUA chromatin. On the basis of bulk histones, mass spectrometry has pointed out several key PTMs, whose abundance depends on whether embryonic cells are proliferating or not. As a reference point serve H3K4me2/me3 marks, located at transcriptional start sites of active genes. Their abundance is well maintained under HUA treatment, indicating that the number, although not necessarily the identity, of active and poised promoters remains constant between the experimental conditions. In contrast, acetylated H3K9 and H3K27 marks are 3-6 fold diminished in arrested embryos, and recover significantly upon inhibitor washout. In functional terms, acetylation protects K9 and K27 residues from becoming decorated with repressive methyl marks. Specifically, the reduction in H3K27ac suggests a problem with the activation of distal enhancers, which diversify cell-type specific transcriptional programs during differentiation (Smith and Shilatifard, 2014). At the same time, HUA chromatin contains about three-fold elevated levels for transcriptionally repressive H3K9me3, H3K27me3, and H4K20me3. To evaluate this increase properly, one has to consider two aspects. First, based on an experimentally determined lengthening of the cell cycle from about 5 hours at gastrulation to about 12 hours at tailbud stage (Thuret, Auger and Papalopulu, 2015), HUA arrested embryonic cells probably miss only three to four rounds of mitosis. Secondly, the normal abundance of the repressive PTMs lies below 5% (Fig S6), which is sufficient to control all heterochromatic and silenced parts of the genome. Therefore, a near three-fold increase bears a significant regulatory potential, either by enlarging the size of repressed chromatin domains, or by increasing the modification density on sites decorated at threshold levels such as bivalent chromatin domains (Li et al., 2018). Under the assumption that losses of acetyl marks on H3K9 and H3K27 are locally coupled to a gain in methyl marks, this altered histone landscape could decrease transcription of many genes in HUA embryos, which would be needed at higher expression levels under normal conditions. This can be investigated in organoid cultures, which compared to embryos have a more homogenous cellular composition and thus are amenable to RNA- and ChIP-Seq profiling.

Which kind of mechanisms link the cell cycle to the histone modification landscape? Studies following the acquisition of PTMs on newly incorporated histone proteins have revealed that H3K27me2/3, H3K79me1/me2 and H4K20me2/3 reach higher levels in starved cells than in cycling cells. This suggests that their steady state depends on the amount of time spent in G1/G0 (Xu et al., 2011; Zee et al., 2012; Alabert et al., 2015). This rationale may also apply to cells transiently arresting in the G2-phase, as it occurs for instance during neurogenesis in frogs and flies (Sabherwal et al., 2014; Otsuki and Brand, 2019). Although the chromatin of these cells has not been profiled, they are expected to contain maximal levels of the repressive mark H4K20me1 on newly incorporated histones, due to the G2/M-phase specific activity of PR-Set7 (Pesavento et al., 2008). An increase in cell cycle length has been shown to control the nuclear localization of MET2. This histone methyltransferase is responsible for the timely accumulation of H3K9me2, which restricts cellular plasticity during C. elegans embryogenesis (Mutlu et al., 2019). While these examples illustrate cell cycle phase-specific influences on histone modifying enzymes, other mechanisms directly influence the inheritance of histone PTMs during S-phase. Most modifications appear to follow the simple paradigm that new histones are modified until they become identical to the old ones, typically in one cell cycle round. In contrast, the modifications H3K9me3, H3K27me3 and probably H4K20me3 (our work here) are propagated with continuous, but much slower kinetics on both old and new histones to restore the parental density (Alabert et al., 2015). While the mechanisms for these kinetic differences have not been resolved, they suggest that cell proliferation is rate-limiting for the maintenance of some histone PTMs. Data from the adult mouse gut strongly support this notion. Intestinal stem cells with a low proliferative rate contain higher levels of H3K27me3 than their more proliferative descendants, i.e. transitory amplifying cells (Jadhav et al., 2020). Furthermore, the conditional knockout of the PRC2 subunit EED in the intestine reactivates cell proliferation and causes derepression of genes proportional to the number of preceding cell divisions. Although most of this data rests on indirect evidence, it suggests a threshold level for H3K27me3, below which transcriptional repression is lost. These findings led the authors to conclude that replicational dilution is the major cause of H3K27me3 removal in mammalian cells (Jadhav et al., 2020). Such a simple model may not be generally applicable to all modifications. Not only there are distinct modes of propagation (fast and slow), but also the combination of different histone marks matter. In the so-called domain model of histone propagation, preexisting methyl marks on K27 and K36 of the histone H3 tail mutually antagonize their respective methylation rates, such that the site of incorporation of new histones determines their ultimate methylation state. This mechanism is thought to enhance the stability of epigenetic states (Alabert et al., 2020). Based on such findings and our data presented here, we expect a highly diversified impact of the cell cycle on the epigenetic landscape.

Finally, does the altered chromatin landscape affect embryonic development? Our analysis allows only to draw indirect conclusions, since we cannot determine the individual distribution of histone modifications in the different experimental conditions due the cellular heterogeneity of the embryo. However, on the way from the tailbud stage to the phylotypic stage (NF28-31), when the vertebrate body plan becomes apparent, we have noticed several differences between control and HUA embryos, which deserve to be discussed.

While gene-expression occurs generally in synchrony between the two conditions (Fig S1), HUA embryos fail to differentiate several externally visible structures. These include: i) melanocytes, i.e. descendants of the cranial neural crest, unambiguously identified by melanin production; ii) a translucent fin, which forms on the dorsal midline along the main body axis; iii) a post-anal tail. The absence of cell proliferation does not explain these phenotypes per se, since related organs are formed, as indicated by the small cluster of rx1-positive cells that is visible in most HUA embryos (Fig 3C) and defines the eye cup primordium. This suggests that either the inducing or the responding cells are compromised in HUA arrested embryos. The upregulation of repressive histone marks coupled with the loss in H3K9ac/H3K27ac marks would fit a scenario, in which genes required in retina have not become activated to sufficient levels. A similar case can be made for the tail, which grows out from two spots of mesendodermal cells, located left and right of Spemann’s organizer (Beck and Slack, 1998). Cells at the tip of the tail maintain expression of the transcription factor genes brachyury (xbra) and foxd5a. Permanently arrested (HUA) embryos have lost the expression of these genes in the tail primordium, unlike transiently arrested (HUAwo) embryos, who maintain their expression and form a proper tail (Fig 3 and 5; Table S1). The tail represents a paradigmatic example, where gene expression could be compromised by changes in histone PTMs in response to cell cycle arrest. These hypotheses can be addressed and deserve further experimentation in the future.

In summary, we propose that in vertebrate embryos the duration of proliferation in cell lineages, and the frequency of mitosis, as a function of cell cycle length, can control the abundance of histone modifications via replicational dilution. As we have argued here, although not proven, this might be sufficient to control developmental decisions. In principle, PTMs with slow biological rate constants are predicted to be preferentially sensitive to this mechanism. On the other hand, cell populations may experience a particularly strong dilution effect, when they switch between resting and proliferating states. While this cell behavior occurs frequently in development (Hall et al., 2010; Buchholz, 2015; Rankin et al., 2015; Thuret, Auger and Papalopulu, 2015), a universal impact for replicational dilution on the epigenetic landscape is expected in progenitor and stem cell populations.

## Materials and Methods

### Ethics statement

Xenopus experiments adhere to the Protocol on the Protection and Welfare of Animals and are approved by the local Animal Care Authorities.

### Embryos handling and HUA treatment

Xenopus laevis eggs were collected, in vitro fertilized and handled as described in “Early Development of Xenopus Laevis: A Laboratory Manual” (Sive, Grainger and Harland, 2000). Embryos were staged accordingly to Nieukoop and Faber (Nieuwkoop and Faber, 1994). When they reached the desired stage, the embryos were immersed into a solution with two DNA synthesis inhibitors (HUA): 20 mM hydroxyurea and 150 μM aphidicolin (made from a frozen stock at 10 mg/ml in dimethyl sulfoxide, DMSO) in 0.1x MBS solution. The first inhibitor Hydroxyurea blocks ribonucleotide diphosphate reductase, an enzyme that catalyzes the reductive conversion of ribonucleotides to deoxyribonucleotides, a crucial step in the biosynthesis of DNA. The second inhibitor Aphidicolin blocks eukaryotic DNA Pol-alpha. To establish a full and irreversible cell cycle block HUA treatment takes 4-6h, if kept in the solution continuously (Harris and Hartenstein, 1991). Within this time period embryos develop from NF10.5 to NF13, if cultivated at 16C. Controls in all cases were embryos from the same batch that were treated identically, except they were pipetted into 2% DMSO in 0.1x MBS solution (Mock). HUA- and Mock-treated embryos were kept continuously in the HUA and Mock solutions respectively, while in the HUAwo condition the embryos were transiently incubated in the HUA from NF10.5 until NF13, and then moved in the Mock until the end point of the analysis, NF32.

### RNA in situ hybridization and immunocytochemistry

Whole-mount RNA in situ hybridization was performed as described in “Early Development of Xenopus Laevis: A Laboratory Manual” (Sive, Grainger and Harland, 2000). For immunocytochemistry anti-H3S10Ph antibody (1:500, Active Motif) and anti-mouse alkaline phosphatase-conjugated (1:2000, Chemicon) secondary antibody was used. Embryos were photographed with a Leica M205FA stereomicroscope. Signal from H3S10Ph+ cells was counted using ImageJ software. For confocal imaging anti-H3S10Ph antibody (1:500, Active Motif), anti-x-β-catenin antibody (1:100, Elizabeth Kremmer) and DAPI were used. Embryos were photographed with a Leica TCS SP5II confocal microscope, using Z-stack montage function. Z-stack depth – 30 micron.

### Nuclear histone extraction

Around 50 to 200 embryos developed to desired stages (NF13, NF18, NF25 NF32) were harvested and washed with 110 mM KCl, 50 mM Tris/HCl (pH 7.4 at 23uC), 5 mM MgCl2, 0.1 mM spermine, 0.1 mM EDTA, 2 mM DTT, 0.4 mM PMSF, 10 mM Na-butyrate. Nuclei of the embryos were prepared by centrifugation with 2600xg, 10 min (3–18, Sigma) after homogenization by a 5 ml glass-glass douncer (Braun, Melsungen). The nuclear pellets of Xenopus embryos were resuspended in 1 ml 0.4 M HCl, incubated on a rotating wheel overnight and dialysed against 3l of 0.1 M acetic acid/1 mM DTT. The dialysed histone solution was vacuum-dried in a Concentrator Plus (Eppendorf) and stored at −20C.

Each developmental stage in the Experiment Type A is represented by three biological replicates; one biological replicate for the Experiment Type B. Each biological replicate derives from a different mating pair.

### Histone acid extraction and sample preparation

The pellet from the nuclear histone extraction was dissolved in an appropriate amount of Laemmli Buffer to reach 1.37e6 nuclei/μl in each sample. 15μL were loaded on an 8-16% gradient SDS-PAGE gel (SERVA Lot V140115-1) and stained with Coomassie Blue to visualize the histone bands. Histone bands were excised and propionylated as described before (Villar-Garea et al., 2012). Isotope heavy-labelled peptides (product of JPT company) 500fmol each were added in the samples before in-gel trypsin digestion. Digested peptides were sequentially desalted using C18 Stagetips (3M Empore) and porous carbon material (TipTop Carbon, Glygen) as described elsewhere (Rappsilber, Mann and Ishihama, 2007) and resuspended in 15μl of 0.1% FA.

### Mass Spectrometry analysis with scheduled PRM method

To study histone post-translational modifications (PTMs) during Xenopus development, we extracted histones from HUA treated and control cohorts of embryos from the 4 developmental stages (Fig S7). An absolute and relative abundance of the histone PTMs were measured on the QexactiveHF LC-MS/MS.

As an internal and inter-sample control a library consisting of isotopically labelled peptides (R10s) was used (Table S2) to normalize for ionization differences between peptides. The R10 peptides were mixed in the library at equimolar concentration and the mix was added to each analyzed sample. In total, the library consists of 65 peptides, 60 of them represent different histone H3 modification states, 6 — histone H4 modification states. Considering only confirmed methylation and acetylation modifications on H3 and H4 histone tails, the library covers 76% of the histone H3 and 50% of the histone H4 modification states.

In order to identify and measure the abundance of the histone PTMs we used a parallel reaction monitoring method (PRM) (Liebler and Zimmerman, 2013). It allows to isolate a target peptide ion based on the mass and retention time window, fragment it and analyze the masses of all fragment ions simultaneously (Fig. S3). Therefore, we could distinguish and identify peaks of isobaric peptides.

The mass spectrometer was operated in a scheduled PRM mode to identify and quantify specific fragment ions of N-terminal peptides histone proteins. In this mode, the mass spectrometer automatically switches between one survey scan and 9 MS/MS acquisitions of the m/z values described in the inclusion list containing the precursor ions, modifications and fragmentation conditions (Table S3). Survey full scan MS spectra (from m/z 270-730) were acquired with resolution 60,000 at m/z 400 (AGC target of 3×10^6). PRM spectra were acquired with resolution 30,000 to a target value of 2×10^5, maximum IT 60 ms, isolation window 0.7 m/z and fragmented at 27% or 30% normalized collision energy. Typical mass spectrometric conditions were: spray voltage, 1.5 kV; no sheath and auxiliary gas flow; heated capillary temperature, 250°C.

### Histone PTM quantification

Data analysis was performed with the Skyline (version 3.7) (MacLean et al., 2010) by using doubly and triply charged peptide masses for extracted ion chromatograms (XICs). Selection of respective peaks was identified based on the retention time and fragmentation spectra of the spiked in heavy-labelled peptides. Integrated peak values (Total Area MS1) were exported as csv. file for further calculations. Total area MS1 from endogenous peptides was normalized to the respective area of heavy-labelled peptides. The sum of all normalized total area MS1 values of the same isotopically modified peptide in one sample resembled the amount of total peptide. The relative abundance of an observed modified peptide was calculated as percentage of the overall peptide.

### RNA extraction and qPCR sample preparation

Total RNA of 10 embryos was extracted using Trizol (Ambion) and phenol/chloroform. The RNA was precipitated with 70% Isopropanol and cleaned using the RNeasy Cleanup Kit (Qiagen) including DNAse I-on-column digestion. For qPCR analysis 1 μg of total RNA was transcribed with the DyNAmo CDNA Synthesis Kit (Bioenzym). For qPCR 5–20 ng cDNA was mixed with the Fast SYBR Green Master mix (Applied Biosystems) and amplified with a Light-cycler (Roche). Primer sequences are given in (Table S4).

### Statistical analysis

For embryonic quantitative analysis (morphological phenotype, qRT/PCR, quantitative MS analysis) SEM are displayed and the statistical analysis was performed using two-tailed, paired Student’s t-test. For boxplots, “ggplot2” R package was used. Principal Component Analyses (PCA) were performed using R, without scaling ().

### Heatmap Generation

Mass spectrometry intensity values of the spiketides were log2-transformed and missing values imputed using two nearest neighbours (library “knnImputation” in R). Endogenous peptides intensities were log2-transformed after adding 1 to all values. For heatmap display (Fig 4A) the normalized log ratios were quantile normalized across all samples and subsequently average per condition (n=3 biological replicates per stage per condition). The values were scaled row-wise per peptide and hierarchically clustered using the “complete” method on Euclidean distances.

## Acknowledgements

We express our gratitude to Edith Mentele and Barbara Hölscher for their expert technical support in RNA in situ hybridization and immunocytochemical staining. We thank Lea Schuh and Carsten Marr for their comments. This work was funded by the Deutsche Forschungsgemeinschaft (DFG, German Research Foundation) – Project-ID 213249687 – SFB 1064.

---

**Supplementary Figure 1.**
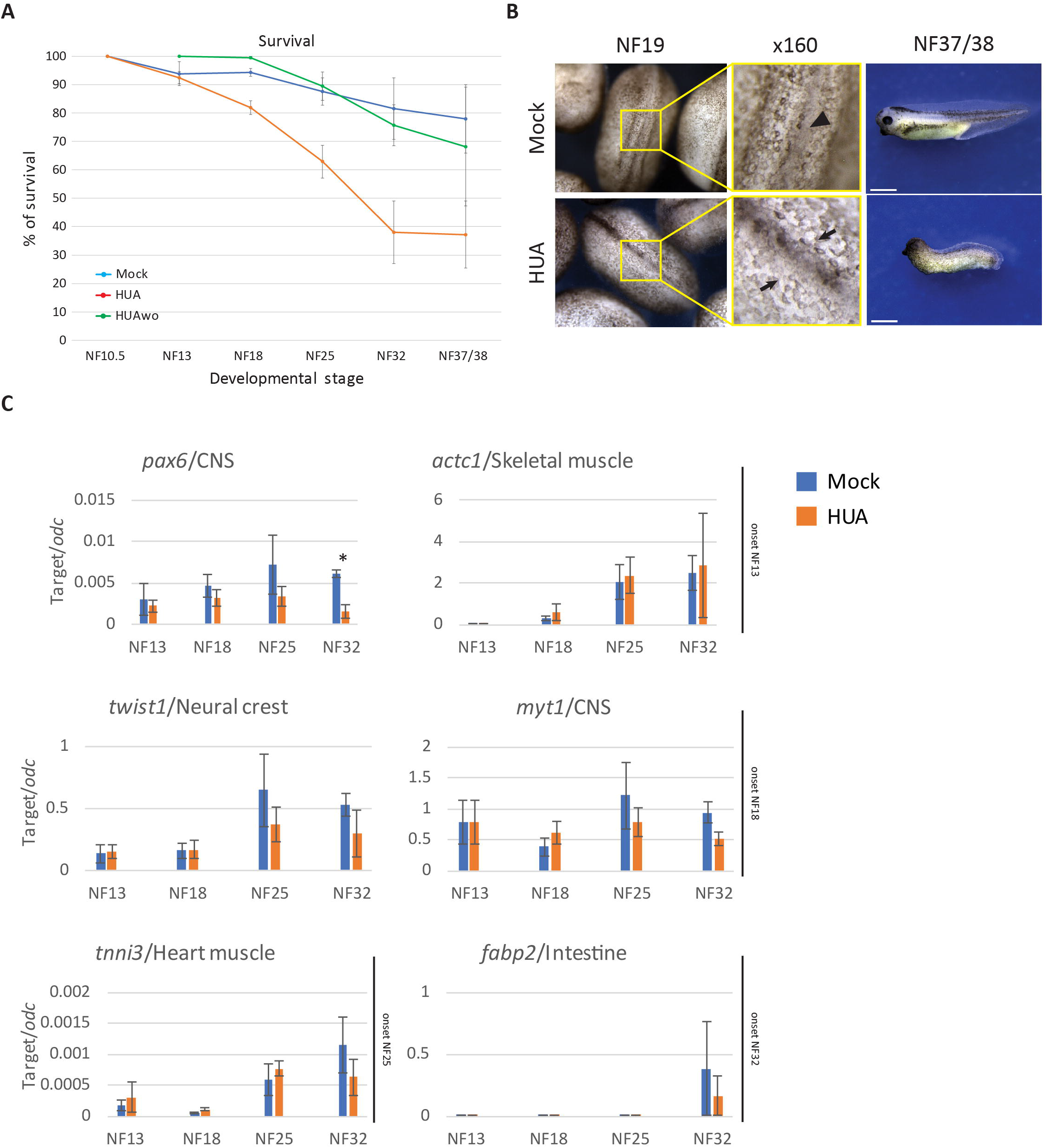
HUA treatment reduces survival and impacts morphogenesis. A) Embryonic survival curves under Mock, HUA or HUAwo condition. Data from n>3 biological replicates/condition; mean ± s.e.m. B) The first morphological effect of HUA treatment is apparent at stage NF19 as a delay in neural tube closure (in mock, black arrowhead points to the dorsal midline; in HUA, two black arrows point to separate neural folds). Under higher magnification, HUA embryos contain larger cells. After hatching (stage NF37/38), HUA treated embryos lack tails, have reduced eyes, malformed fins and are largely deficient in melanocytes. C) Comparison of temporal expression profiles for selected marker genes in Mock and HUA conditions by qRT/PCR, normalized to odc mRNA. Genes are grouped according to their activation time point. N=3 biological replicates/condition; mean ± s.e.m. Significant difference was detected only in case of pax6 expression level at NF32 stage (Student’s t-test [twotailed, paired]; * p<0.05). No other significant differences were detected between the two conditions, indicating a proportionate gene expression pattern, consistent with the increasing developmental age.

**Supplementary Figure 2.**
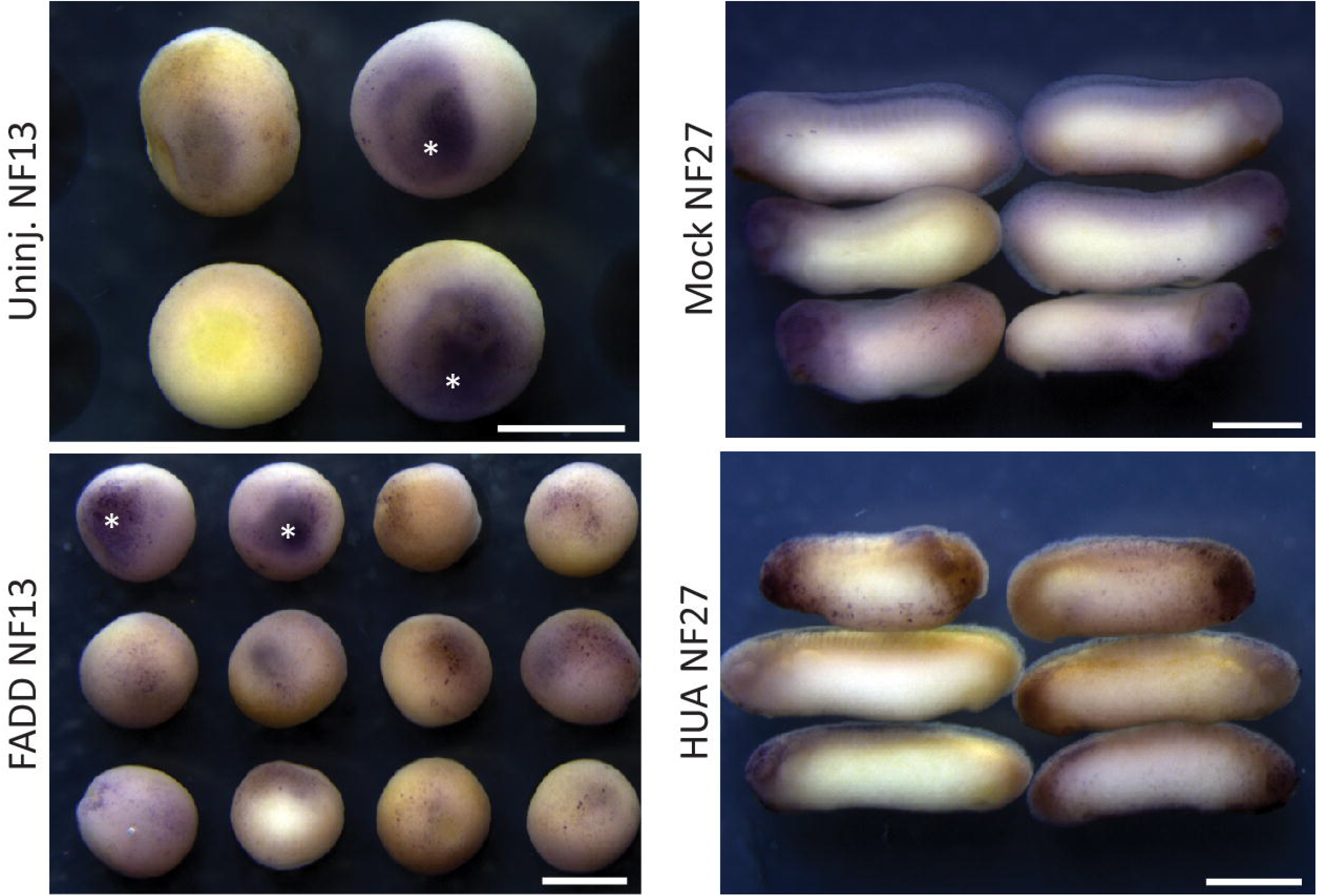
Apoptosis in HUA and Mock treated embryos. Wild type embryos were injected with FADD apoptosis inducing plasmid in one blastomere at 4-cell stage as a positive control. FADD injected and uninjected wild type embryos together with Mock and HUA embryos were ICC stained against activated Caspase-3. Signal from cas-3 staining can be observed on the skin of the embryos as small blue dots. White asterisks indicate blastocoel background staining, scale bars: 1mm. HUA embryos do not demonstrate an increased level of apoptosis compared to Mock siblings.

**Supplementary Figure 3.**
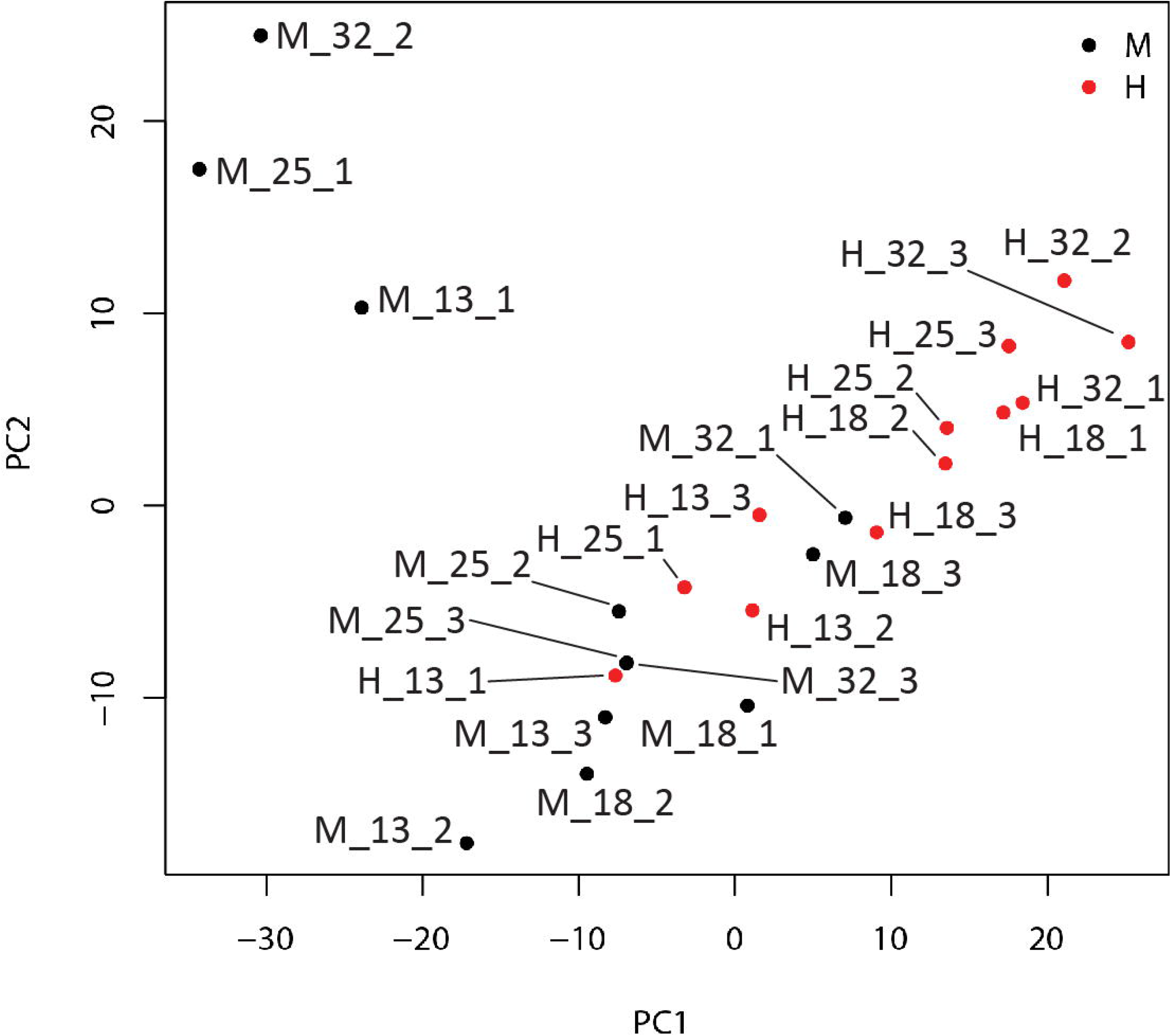
Principal component analysis for Mock and HUA-treated HPMs. Each data point represents 65 modification states, measured by LC-MS/MS in PRM mode, with absolute abundances calculated with R10 spiketide normalization. Mock and HUA data sets are partially separated, with younger HUA samples intermingling with older Mock samples.

**Figure 4:**
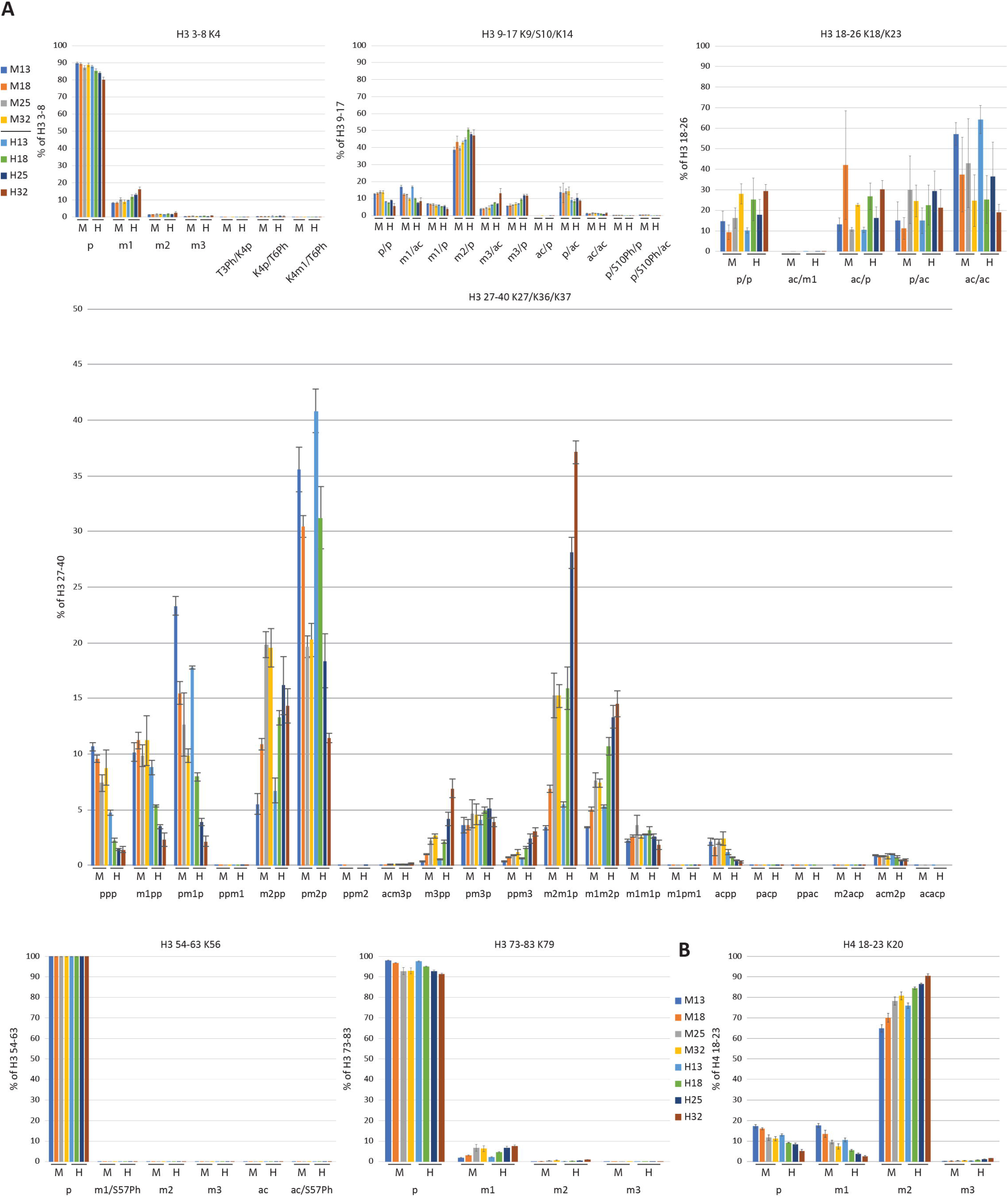
Relative histone PTM abundances in HUA and Mock treated embryos. A) Individual relative histone PTM distribution for histone H3. Data is first normalized to R10 spiketide signals, then added up to 100% for all modification states measured for each specific tryptic peptide, from which the relative contribution of each state is then calculated. B) Individual relative histone PTM distribution for histone H4. N=3 biological replicates/condition; mean ± s.e.m.

**Supplementary Figure 5.**
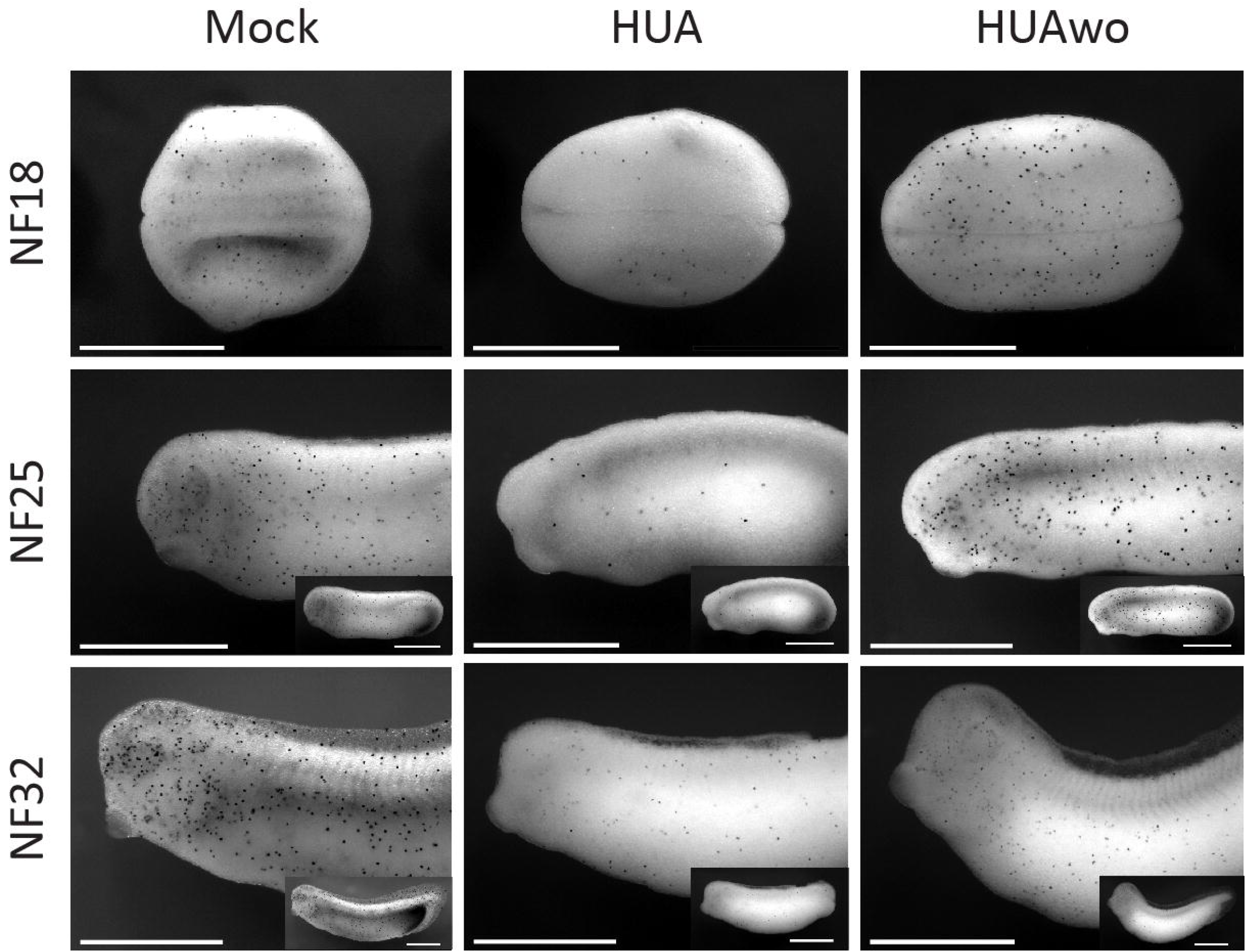
Recovery of mitotic activity in HUAwo embryos. Immunocytochemical staining (ICC) for the mitotic histone mark H3S10Ph at indicated stages. Mitotic cells are marked by black dots. Elongated, older embryos are recorded as anterior halves, i.e. at same magnification as younger stages, and in whole mount views as inserts. Scale bars: 1mm. N=3 biological replicates/condition.

**Supplementary Figure 6.**
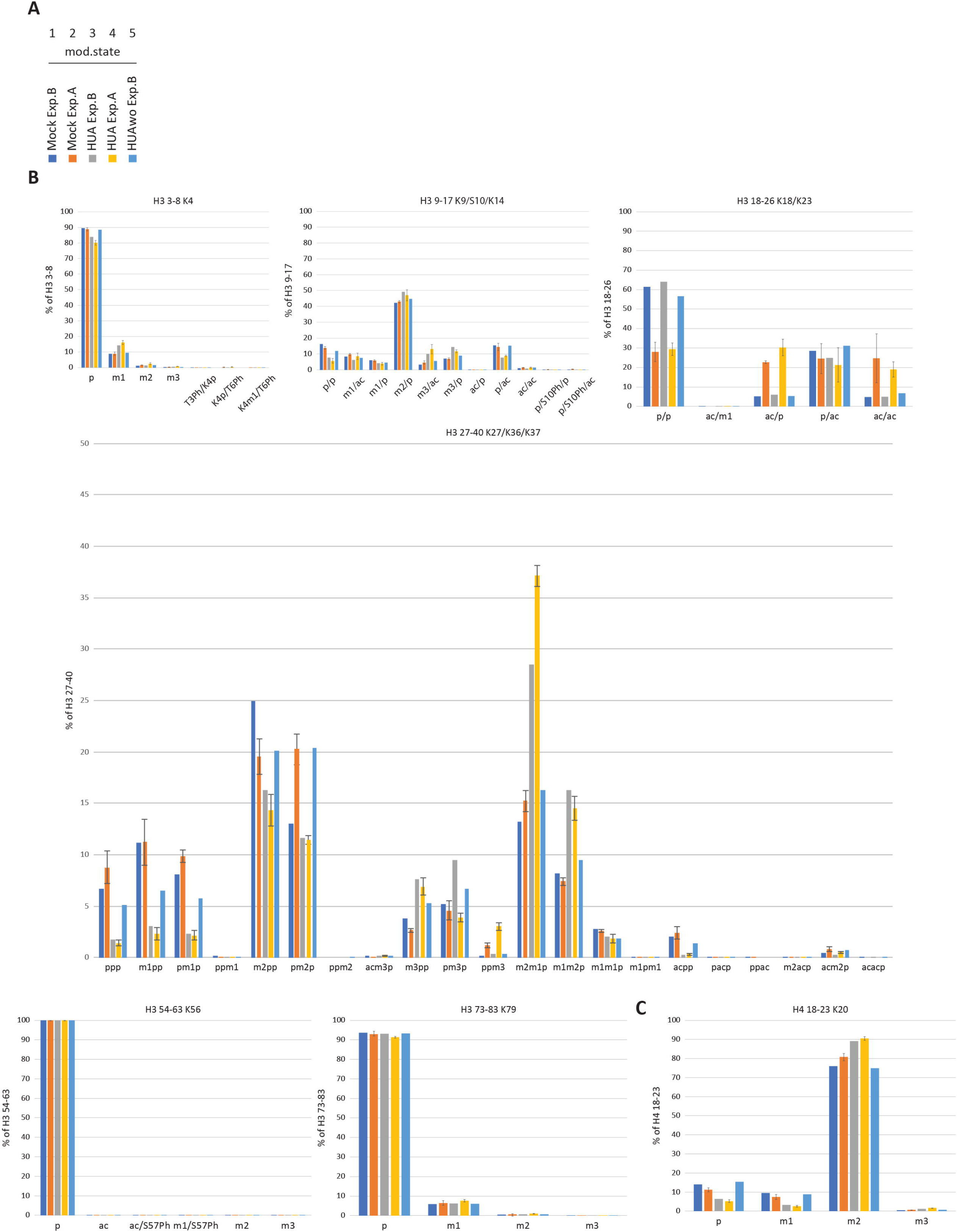
Comparison of relative histone PTM abundances from Experimental series type A and B. Relative PTM abundances were calculated as described in Materials and Methods. A) Color coding of sample types. Panels B and C: Individual relative histone PTM distributions for histone H3 and H4, respectively.

**Supplementary Figure 7.**
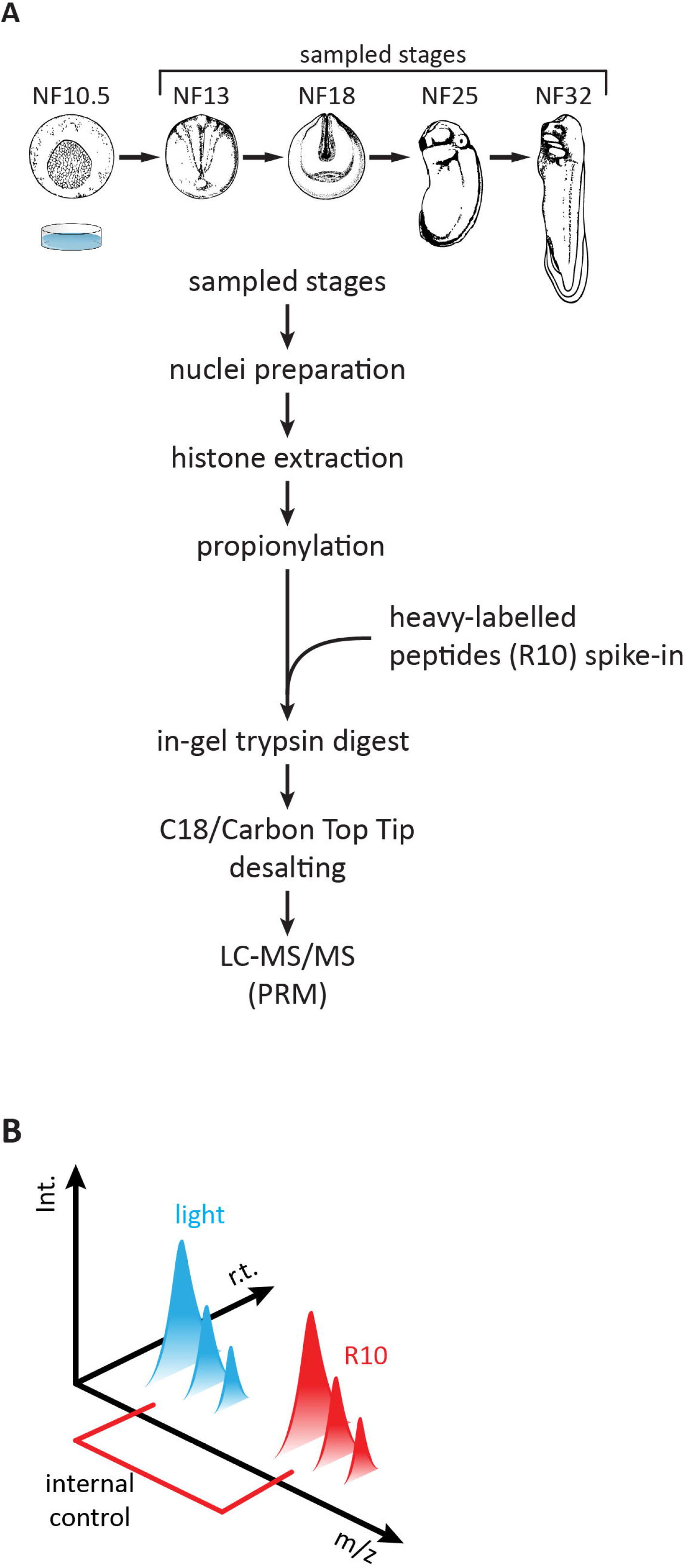
Absolute quantification of histone post-translational modification states by LC-MS/MS using scheduled PRM method. A) Pipeline of mass spectrometry analysis of histone modifications from Xenopus laevis. Bulk histones are isolated from purified nuclei of embryos from four sampled stages (see Fig 1) by acidic extraction and SDS-PAGE. Propionylation blocks all endogenously unmodified and monomethylated Lysine residues from being cleaved in the subsequent trypsin digest, thereby creating an optimized peptide pool for Mass Spec analysis. After propionylation, but before trypsin digest, we add to each sample a so-called R10 library (Table S2), which consists of isotopically heavy-labelled Arginine peptides (R10). The individual R10 peptides are mixed in equimolar concentration and mimic 65 histone H3 and H4 modification states. These isotopically heavy-labelled peptides serve as an internal and inter-sample control, allowing to minimize technical variations and to quantitate abundances of histone modification states on the absolute scale. B) Representation of the R10 spike-in peptide control. Each of the analyzed endogenous histone modification states has a synthetized R10 peptide analogue. Due to the same chemical properties, endogenous tryptic peptides and their R10 spiketide analogues elute at the same retention time (r.t.); however, they can be distinguished based on the mass to charge (m/z) ratio. Additionally, R10 spiketides help with peak identification based on retention time and detail fragmentation spectra for isobaric peptides.

